# Aridity drives coordinated trait shifts but not decreased trait variance across the geographic range of eight Australian trees

**DOI:** 10.1101/2020.02.03.932715

**Authors:** Leander DL Anderegg, Xingwen Loy, Ian P. Markham, Christina M Elmer, Mark J Hovenden, Janneke Hille Ris Lambers, Margaret M Mayfield

## Abstract

**Context:** Large intraspecific functional trait variation strongly impacts many aspects of natural communities and ecosystems, yet is inconsistent across traits and species.

**Approach:** We measured within-species variation in leaf mass per area (LMA), leaf dry matter content (LDMC), branch wood density (WD), and allocation to stem area vs. leaf area in branches (branch Huber value, HV) across the aridity range of seven Australian eucalypts and an *Acacia* species to explore how traits and their variances change with aridity.

**Results and Conclusions:** Within-species, we found consistent increases in LMA, LDMC and WD, and HV with increasing aridity, resulting in consistent trait coordination across tissues. However, this coordination only emerged across sites with large climate differences. Unlike trait means, patterns of trait variance with aridity were mixed across populations and species and showed limited support for constrained trait variation in dryer populations or more xeric species.

**Synthesis:** Our results highlight that climate can drive consistent within-species trait patterns, but that these patterns might often be obscured by the complex nature of morphological traits and sampling incomplete species ranges or sampling confounded stress gradients.

## Introduction

Land plants exhibit astounding variation in both form and physiological function. The identification of ‘functional traits’ as easily measured plant attributes that are proxies for plant physiological function and performance has spurred the rise of the field of ‘plant functional ecology’ and revealed some of the key causes and consequences of plant functional diversity (Mooney, Ferrar & Slatyer 1978; Field 1988; Reich, Walters & ellsworth 1997; Díaz *et al.* 2016; Ma *et al.* 2018). In particular, across-species studies of plant traits have revealed global ‘trait spectra’ or ‘trait syndromes’—correlations between different plant traits indicative of coordination across various aspects of plant physiology—that both illuminate trade-offs shaping plant evolution and provide powerful tools for community and ecosystem ecological studies (Wright *et al.* 2004; Reich *et al.* 2014; Ma *et al.* 2018).

While functional ecology has largely been built on trait patterns among species, our understanding of trait variation and trait coordination within individual species remains more limited. Ecologists increasingly recognize that within-species trait variation can be a large fraction of total trait variation (Albert *et al.* 2010b; Siefert *et al.* 2015), and that within-species trait variation has large consequences for ecological and evolutionary processes (Lepš *et al.* 2011; Laforest-Lapointe, Martinez-Vilalta & Retana 2014), particularly the potential for species to adapt to climate change. Within-species variation in functional traits linked to stress tolerance has been increasingly used to predict plant responses to global change (Martinez-Vilalta *et al.* 2009; Laforest-Lapointe *et al.* 2014; Anderegg & HilleRisLambers 2015; la Mata, Hood & Sala 2017). Even as our appreciation of the importance of intra-specific variation grows, a mounting body of perplexing results reveals the limits to our understanding of within-species trait variation. For example, within-species trait responses to environmental gradients have sometimes defied generalization by proving highly trait-specific and species-specific (e.g. Schulze *et al.* 1998; Albert *et al.* 2010b; a; Vilà-Cabrera, Martinez-Vilalta & Retana 2015; Rosas *et al.* 2019)1, and sometimes even study specific (e.g. Martinez-Vilalta *et al.* 2009; Laforest-Lapointe *et al.* 2014).

The between-species trait-trait coordination that underpins theory about trait spectra does not necessarily hold within individual species (Messier *et al.* 2017; Anderegg *et al.* 2018; Messier *et al.* 2018). For instance, a recent analysis of intra-specific trait coordination in saplings of temperate tree species found that essentially none of the canonical trait relationships behind three classic theories of trait coordination held across populations within species (Messier *et al.* 2018). In another example, strong between-species trait-by-environment relationships and trait coordination across an aridity gradient in northern Spain generally failed to emerge across populations within individual species along the same gradient (Rosas *et al.* 2019). Indeed, some important trait-trait relationships can even reverse direction within-versus between-species (Anderegg *et al.* 2018). This contrasting within-verse between-species trait coordination suggests that classical explanations of trait integration do not necessarily hold within-species, limiting their applicability for predicting species’ functional responses to climate change.

Not only are within-species trait-by-environment relationships and trait coordination uncertain, patterns of trait variances within-species remain poorly understood. The study of trait variances (rather than trait means) has a long history in community ecological and evolutionary studies, yet trait variances have often been overlooked in the ecophysiological literature. The amount of local adaptation and phenotypic plasticity, the two sources of within-species variation, are, however, key constraints on species responses to climate change (Richter *et al.* 2011; Chevin, Collins & Lefèvre 2012; Alberto *et al.* 2013; Franks, Weber & Aitken 2014; Valladares *et al.* 2014) Predicting plant responses to a shifting environment requires an improved understanding of total trait variation in different species and between populations of the same species (Molina-Montenegro & Naya 2012; Lemke, Kolb & Diekmann 2012; Siefert *et al.* 2015).

Increased leaf robustness quantified by Leaf Mass per Area (LMA) and Leaf Dry Matter Content (LDMC) and stem robustness quantified by Wood Density (WD) are often associated with xeric environments because their variation is partly driven by anatomical adjustments that allow plants to maintain hydraulic function under increasingly negative xylem pressures (Schulze *et al.* 1998; Niinemets 1999; Schulze *et al.* 2006; Chave *et al.* 2009; Poorter *et al.* 2009; John *et al.* 2017). The ratio of stem sapwood area to leaf area or Huber value (HV) reflects the balance of hydraulic supply (sapwood area) relative to hydraulic demand (leaf area), with high HV typically indicating increased hydraulic efficiency and thus increased drought avoidance (Tyree & Ewers 1991; Mencuccini & Grace 1995). In single-species studies, different tree species have been found to adjust at least one of these traits with water availability (e.g.. (Martinez-Vilalta *et al.* 2009; Anderegg & HilleRisLambers 2015).

Here, we examine within-species variation in leaf and stem robustness and allocation within closely related tree species across large gradients in water availability in the absence of major confounding environmental stressors, notably freezing. We present a controlled test of predictions about intraspecific trait variation across nested scales of organization, focusing on trait variation across aridity gradients in Western Australia and Tasmania. Further, we minimize differences in species life history by holding phylogenetic history relatively constant for seven core species (‘eucalpyts’ from the closely related *Eucalyptus* and *Corymbia* genera), compared to an addition regionally co-occurring but unrelated species (*Acacia acuminata*).

The specific questions we ask are:

1. Do leaf and stem tissues, and leaf vs stem allocation show consistent relationships with water availability across a species’ range, regardless of how mesic/xeric the species’ dry range edge is?
2. Do species consistently show coordination between leaf and stem robustness, and leaf to stem allocation, and if so at what scale does this coordination emerge?
3. Is the amount of within-species variation in leaf and stem traits more constrained in dry sites (both within species across sites and across sister species with different aridity niches)?

Given their association with drought resistant phenotypes, we expected mean leaf robustness (measured by LMA and LDMC), stem robustness (measured by WD) and allocation to stems compared to leaves (HV) to increase with aridity, resulting in coordinated trait changes across tissues. In addition to expectations about mean trait values, theories on directional selection and environmental filtering led to the prediction that trait variation within and among species should decrease as the climate becomes harsher (Falconer 1989; Kraft *et al.* 2014). Assuming ongoing directional selection towards higher leaf and stem robustness and allocation to stems over leaves with increasing water limitation and a limit to genetic variation, within-species variation in these traits should decrease in more xeric species and within-population variation should decrease in higher aridity populations within a species. A previous metanalysis did not find decreased within-species trait variances in more xeric species (Siefert *et al.* 2015), but we hypothesized that the pattern would emerge in a phylogenetically controlled comparison between closely related species.

## Methods

### Study site

We collected trait data along two temperate aridity gradients (Figure S1), one in southwest Western Australia (sampled November 2014) and one in Tasmania (sampled February 2016). Along each gradient, we identified three to four dominant eucalypt tree species (from the *Eucalyptus* or *Corymbia* genera of the Myrtaceae family) that are easily identified in the field and do not widely form cryptic hybrids or have notable subspecies within the sampled regions. In Western Australia, we sampled *Eucalyptus marginata* Donn ex Sm., *Eucalyptus salmonophloia* F.Muell., and *Corymbia calophylla* (Lindl.) K.D. Hill & L.A.S. Johnson. We also opportunistically sampled the non-eucalypt *Acacia acuminata* Benth., which broadly co-occurs with *E. salmonophloia.* In Tasmania we sampled *Eucalyptus amygdalina* Labill., *Eucalyptus obliqua* L’Hér., *Eucalyptus ovata* Labill., and *Eucalyptus viminalis* subsp. *viminalis* Labill, all of which cover the majority of their global precipitation range within Tasmania. Collectively, sampled sites spanned a mean annual precipitation range of 328 to 1574mm/year (328 to 1189 mm in Western Australia, 584 to 1574 mm in Tasmania) and a moisture deficit (MD, or annual potential evapotranspiration minus annual precipitation) of −755 to 1326 mm deficit/year (−75 to 1326 mm in Western Australia, −755 to 416 mm in Tasmania). Mean annual temperature spanned 8-20°C and elevation ranged from 24-620 m.a.s.l, with no site experiencing significant frost (mean coldest month minimum temperature was greater than 0°C for all sites). Average site climate, soil, tree size and stand basal area (Tasmania only) characteristics can be found in Table S1. Sampled tree size and (where measured in Tasmania) stand Basal Area did not vary strongly with aridity for most species (Table S1). Vegetation in both transects ranged from closed canopy forest to open woodland/savannah. Climate data for sampled plots, including mean annual precipitation (PPT), potential evapotranspiration (PET), and moisture deficit (MD = PET – PPT), were extracted from the Global Aridity and PET Database (Zomer *et al.* 2006), based on the 1960-1990 climate averages of WorldClim (Hijmans *et al.* 2005). An alternative metric of water availability, the Aridity Index (P/PET) was also calculated but found to be almost perfectly collinear with MD at the study plots (Figure S2). Soil properties including soil depth and regolith depth, as well as % sand, silt and clay, total nitrogen by mass, total phosphorus by mass, average water holding capacity, bulk density, and effective cation exchange capacity (averaged over the top 60cm soil depth) were downloaded from the Soil and Landscape Grid of Australia (Grundy *et al.* 2015), using the *slga* R package (O’Brien). Because soil properties were strongly collinear with each other, we performed a principle component analysis (PCA) on the soil variables and represented soil variation in our analyses using the first two principle components (PCs). The first PC explained 67% of total soil variation and was interpreted as general ‘soil fertility’ because it loaded strongly (>0.3) with everything except depth of regolith, depth of soil and water holding capacity. The second PC captured 12% of variation, loaded strongly with water holding capacity and soil depth and was interpreted generally as ‘soil depth’.

### Trait measurement

We measured branch wood density (WD), leaf mass per area (LMA) and leaf dry matter content (LDMC) as metrics of stem and leaf robustness, and terminal branch Huber value (HV), the ratio of sapwood area to leaf area, as a metric of investment in water transport versus light capture. Trait measurements were collected in a nested hierarchical design with four to five sites sampled per species to capture broad climate gradients, three plots per site to capture topographic/edaphic variation, five trees per plot to capture within-population variation, and three samples per tree to capture within-individual variation (Figure S1). For each species, four to five forestry reserves, National Parks, State Forests, Nature Reserves, or Conservation Areas were selected to cover as much of the species’ aridity niche as possible, defined for each species by the moisture deficit (MD, annual potential evapotranspriation minus annual precipitation) of all herbarium specimen locations from the Australia Virtual Herbarium (www.avh.chah.org.au). Edaphic variation within sites was captured by locating three plots that were >500 m but <5 km apart and each containing more than five individuals of the focal species within a 50 m radius. In each of the three plots, we sampled within-population variation by collecting three sun exposed branches from the north side of each of five mature, healthy individuals using pole clippers and pull ropes. Our sample design resulted in 180-225 trait measurements per species.

From each branch, we collected a section ∼8 mm in diameter for WD measurement, and a terminal branch (first order branch collected at the point of branching) for leaf and HV measurements. We selected terminal branches with intact ‘mature’ leaves (i.e. fully expanded, not soft green new growth), though most of the study species flush sporadically throughout the year (Davison & Tay 1989; Heatwole *et al.* 1997) so it was not possible to perfectly control for leaf age. Sampling periods (Nov. in Western Australia and Feb. in Tasmania) avoided large leaf flush events for all species with the exception of *Corymbia calophylla* at two of its five sample sites. Samples were rehydrated in moist ziplock bags in a cooler for at least 12 hours prior to trait measurement (Pérez-Harguindeguy *et al.* 2013). Bark was peeled from branch sections and WD quantified from segments roughly 7 cm in length. WD was weakly related to branch diameter for six species, so diameter was included as a covariate in statistical models of WD for these species.

All leaves subtending the selected terminal branch were collected for measurement of leaf area, LMA and LDMC. Total one-sided leaf area (including petioles) of terminal branch samples was measured with a flatbed scanner and ImageJ image processing software (Schneider, Rasband & Eliceiri 2012). Leaves were then oven dried at 70°C to a constant weight and their dry mass measured. Terminal twig basal diameter was measured just above the swelling at the branch base after gently peeling back bark (except in *A. acuminata*, where bark was difficult to distinguish from woody tissue). For each terminal branch HV, LMA, and LDMC was calculated. Multivariate trait outliers were visually diagnosed and removed (n<10 per trait), as were LMA and LDMC values from still expanding leaves (<10% of measurements).

### Statistics

#### Q1 – Trait-aridity relationships

We tested for significant trait-environment relationships using information-theory based model selection. For each species, we fit linear mixed effects models relating each trait to plot mean annual PPT, PET, MD, soil fertility (soil PC1), soil depth (soil PC2), tree DBH, or stand Basal Area, with plot and tree random intercepts. We then compared the various trait-environment models and a null model (with only a fixed intercept and tree random effect) using Akiake’s Information Criterion (AIC) and selected the model with the lowest AIC. We quantified statistical significance of the most parsimonious model compared to the null model using Likelihood Ratio Tests (LRT). Where a soil variable proved the best trait predictor, we also tested the significance of the best climate model because soil and climate variables were often strongly collinear (Figure S2).

#### Q2 – Trait coordination

We assessed trait-trait covariation using multiple approaches. First, for each species we tested for significant Pearson correlations between tree-level averaged traits for all trait pairs. Next, we assessed the distribution of trait-trait correlations for hierarchically nested data subsets to assess at what level trait coordination emerges. For each trait pair for each species, this involved calculating the Pearson correlations across the replicate branches within each tree, across tree averages in each plot, across plot averages in each site, and across site averages, for all eight sampled species. Lastly, we assessed the dominant mode of trait covariation across all traits and species. We performed a principle component analysis (PCA) on all branch-level trait measurements with complete trait data (1400 branches), and assessed the trait loadings along the first and second PC axes. We then calculated the PC score for all site-averaged trait values, and assessed whether any PC related to site MD across species using a linear mixed effect model including a fixed effect of MD and species random slopes and intercepts.

#### Q3 – Constrained variance at high aridity

We first examined whether more xeric species showed less intraspecific trait variation than mesic species. For each species, we quantified the amount of trait variation at each nested scale for each trait using variance decomposition. For each species and trait, we fit a linear mixed effect model with a fixed intercept and random effects for site, plot, and tree. In this formulation, the random effect variance parameter estimates represent the between-site, between-plot in site, and between-tree in plot variance (respectively), with the residual variance representing variance of samples within tree. To test for across-species patterns, we extracted the variance parameters for each eucalypt species (excluding *Acacia acuminate*) and used linear models to relate species total trait variance (sum of all variance components for a trait) to the species’ driest 90^th^ percentile MD value from occurrences in the AVH database (see above). We also tested whether individual variance components decreased with increasing aridity by fitting linear models relating species variance components to each species’ driest 90^th^ percentile MD plus a fixed effect for variance component (between-site, between-plot, between-tree, or within-tree) and a component-by-MD interaction.

To test for decreasing trait variation with increasing aridity within species (i.e. across populations), we used AIC to determine whether the best trait-aridity mixed effect model (from Q1) for each species and trait was improved by allowing the variance to change as either a power or exponential function of the dominant climate variable, or to assume a different value for each site. If AIC and LRTs suggested that a non-constant variance function improved the trait-climate model, we classified whether the variance increased with aridity, decreased with aridity, or showed variation between sites that was not aridity-related (i.e. the model with different variances per site was the best model). We visualized these within-species variance patterns by plotting the distribution of trait standard deviations within individual trees and within plots.

All analyses were performed in the R statistical environment (Team 2016), version 3.2.4). Mixed effects models were fit using the *lme4* and *lmerTest* packages (Bates *et al.* 2015; Kuznetsova, Brockhoff & Christensen 2016) for fixed variance models, or the *nlme* package for models with more complicated variance structures (Pinheiro *et al.* 2016). Standardized major axis (SMA) regressions were fit using the *lmodel2* package (Legendre 2014). Data available in the Dryad data repository ([insert cite upon publication], https://XXXXXX), and analysis code is available in the Github repository (https://github.com/leanderegg/EucTraits) associated with this paper.

## Results

### Do traits respond to aridity?

For the majority of our examined species, most traits shifted in a way consistent with greater drought resistance (increased WD, LMA, LDMC and HV) in higher aridity plots (Figure 1). All species showed significant trait-by-environment relationships for WD, LMA and LDMC and seven of the eight species showed significant trait-by-environment relationships for HV (Table S1). A measure of aridity (PPT, PET or MD) was the best predictor in 21 of 32 trait-by-environment relationships, soil fertility in 7 of 32 and soil depth in 3 of 32. However, in all but one of the trait-by-environment relationships where soil quality or depth was the best predictor, precipitation was collinear to that soil variable and also a significant, if worse, predictor (Table S1). Precipitation, potential evapotranspiration, moisture deficit and soil fertility were correlated across plots for many, but not all species (Figure S2).

**Figure 1:**
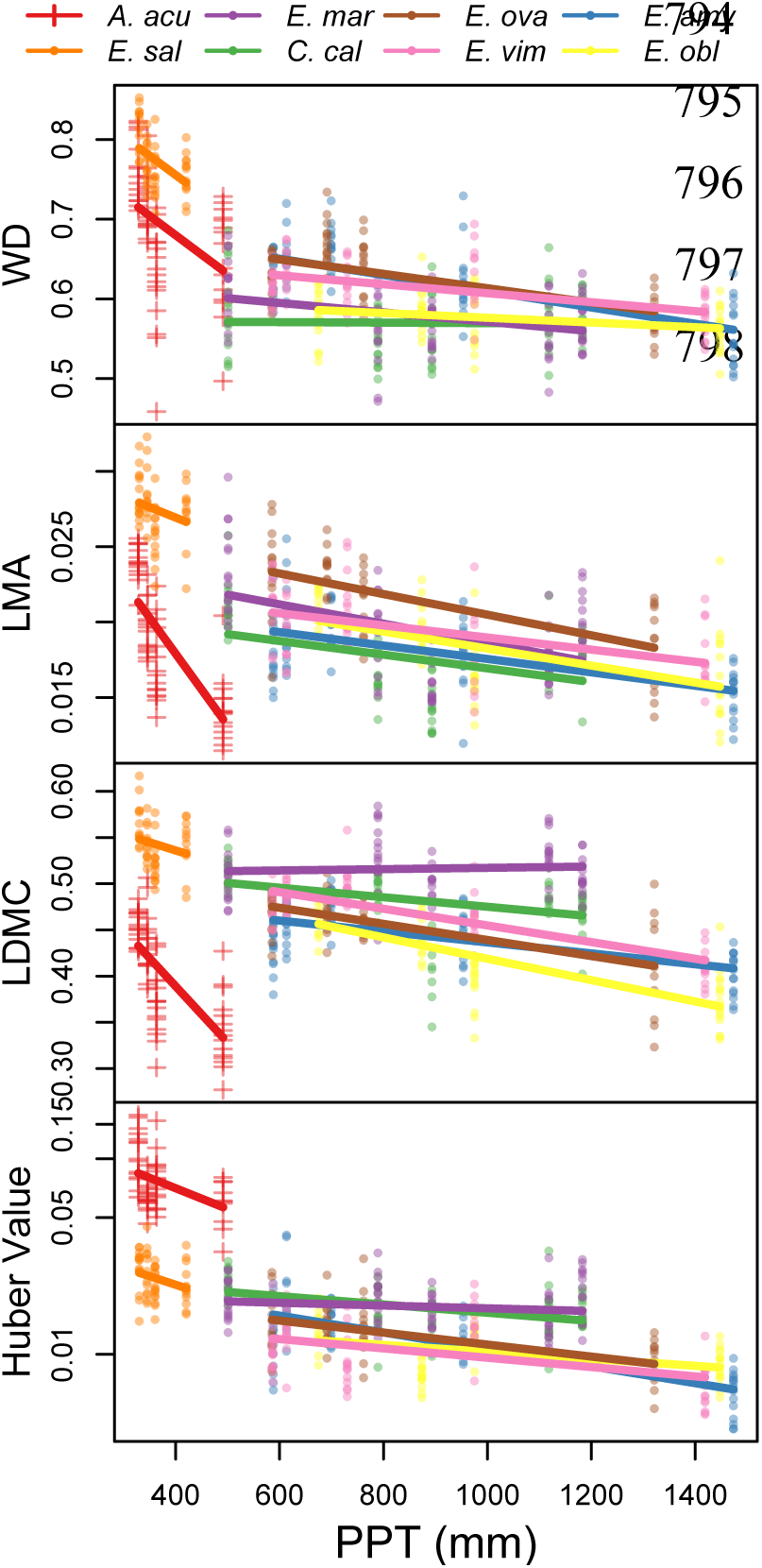
Relationships between four leaf and stem traits and plot mean annual precipitation (PPT) for eight tree species. Points show tree averages. Red crosses show *Acacia acuminata*, the one non-eucalypt species. Species abreviations: *A. acu* – *Acacia acuminata, E. sal* – *Eucalyptus salmonophloia, E. mar* – *E. marginata, C. cal* – *Corymbia calophylla, E. ova* – *E. ovata, E. vim* – *E. viminalis, E. amy* – *E. amygdalina, E. obl* – *E. obliqua*

### Are trait responses coordinated across tissues?

Ubiquitous trait-by-environment relationships resulted in coherent trait coordination across leaf and stem tissue, and coordination between leaf robustness and increased HV within species (Figure 2). However, while consistent and often significant, these within-species trait correlations were typically weak, with the mean within-species trait correlation being <0.5 for all trait pairs except LMA and LDMC. Across tree-level trait averages, the majority of species showed significant correlations between both WD and LMA (mean correlation of 0.33) and WD and LDMC (mean correlation of 0.38; Figure 2a, 2b), though these were typically less strong than the correlations between LMA and LDMC (mean correlation of 0.74; Figure 2c). Both leaf traits were also positively correlated with HV, with mean correlations of −0.44 and −0.32 for LMA and LDMC respectively. However, WD was only significantly correlated with HV in three species. In the seven eucalypts, most species fell in roughly the same trait space, with more trait variation within each species than across species (Figure 2). *Acacia acuminata* showed larger HV, but similar trait correlations to the seven eucalypts (Figure 2d, 2e, 2f).

**Figure 2:**
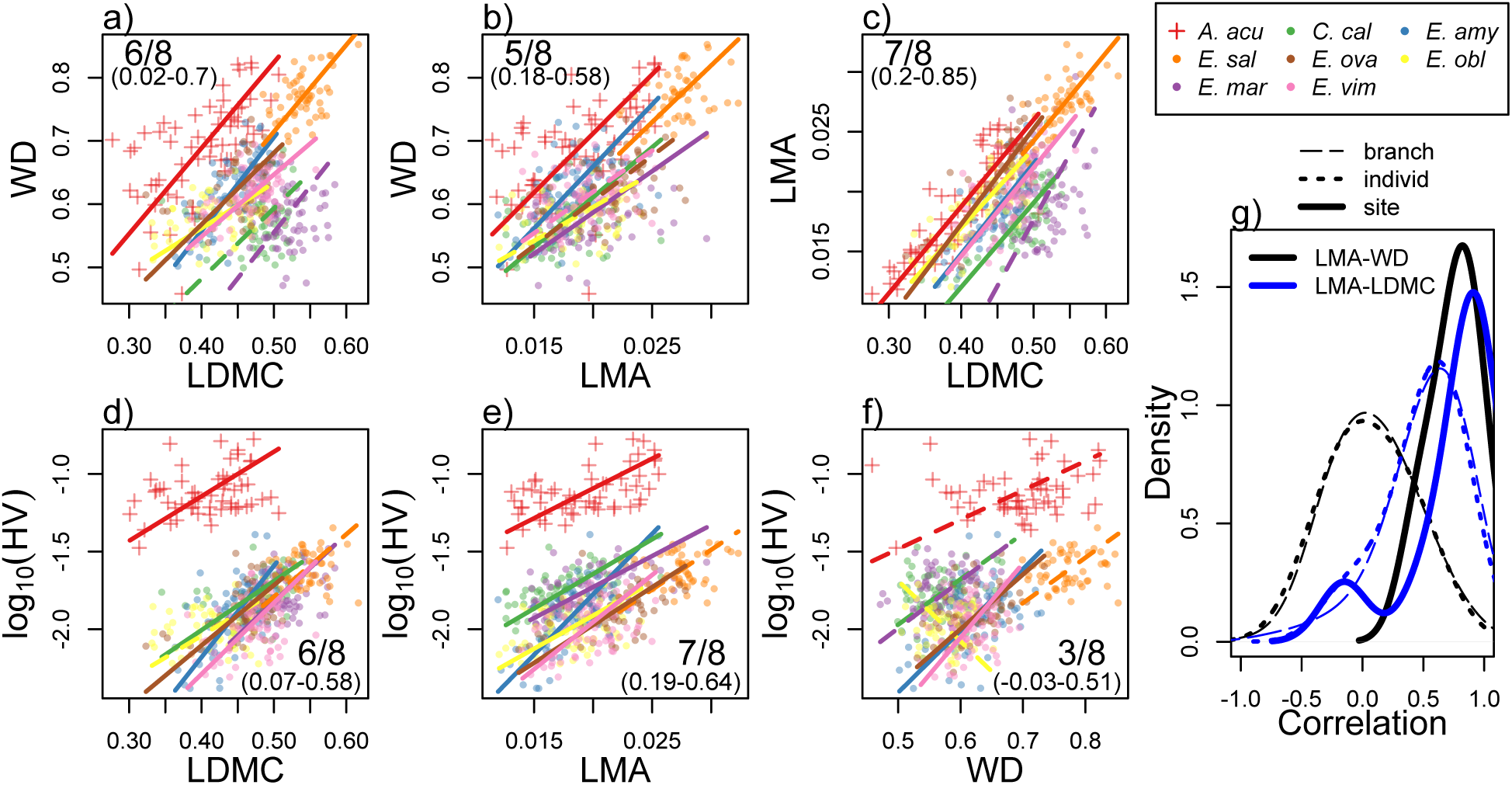
Correlations between leaf and stem traits across the aridity range of eight tree species (a-f). Points show tree average trait values, and lines show Major Axis Regressions (solid lines show significant correlations). Crosses show *A. acuminata*, the one non-eucalypt species. Panel (g) shows the distribution of correlation coefficients across all species for two example trait pairs, LMA vs WD (black) and LMA vs LDMC (blue). Trait correlations typically had a mean near zero across branches or across individuals within a site for all trait pairs except LMA vs LDMC. Species abbreviations as in Fig. 1.

Within-species trait coordination only emerged when comparing traits across the most disparate environments. The distribution of correlation coefficients at smaller spatial scales (e.g. trait-trait correlations across individuals or branches within a plot, correlations across plots or individuals within a site) typically had an interquartile range spanning zero for all trait pairs except LMA-LDMC and HV-LMA(Figure 2g, Table S2). Only when comparing across site mean trait values did the mean within-species correlation differ substantially from zero for most trait pairs (Table S2). This decrease in correlation strength at smaller spatial scales was not purely a result of smaller sampled trait variation, as there was often as much or more trait variation within plots as across sites, and funnel plots did not show strong relationships between correlation strength and sampled trait variance except for the relationship between HV and LMA (Figure S3).

Even though trait coordination only emerged across large aridity gradients, the dominant mode of trait variation in the entire dataset was a coordinated increase in tissue robustness and HV, likely driven by decreasing water availability. In a PCA of the entire branch-level dataset, the first principle component (PC1) explained 53% of the total variance and was loaded reasonably equally with all four traits (Figure 3a). Additionally, for each species the site-level average PC1 score was strongly related to site Moisture Deficit (linear mixed-effects model, p<0.001, marginal R^2^ = 0.53), indicating that the coordinated increase in WD, LMA and LDMC, and HV represented by PC1 was driven by water availability (Figure 3b).

**Figure 3:**
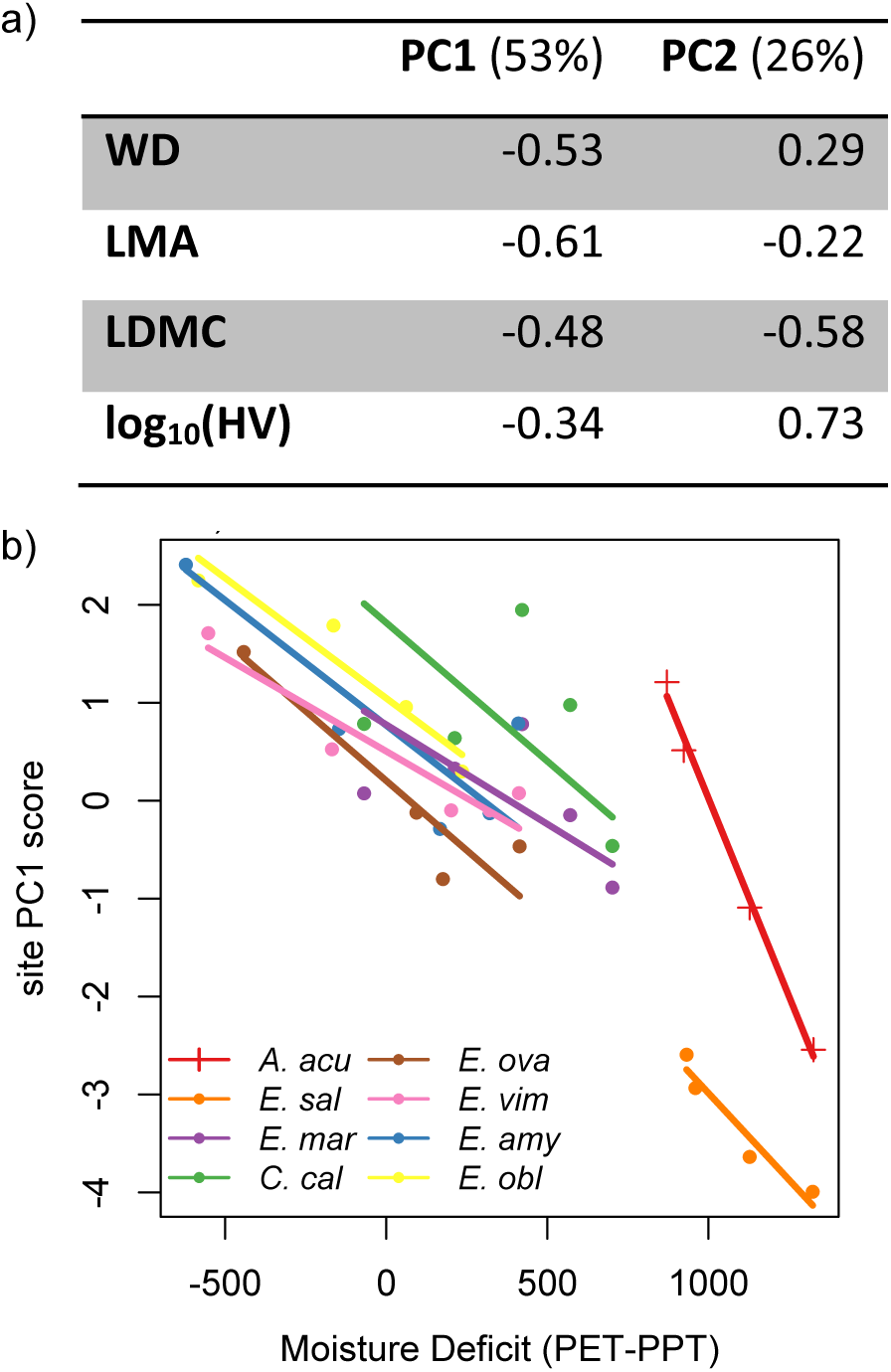
PC loadings of a PCA including all branch measurements (a). Site average trait PC1 scores are strongly related to site moisture availability across eight tree species (b). Species abbreviations as in Fig. 1. PET = potential evapotranspiration, PPT = precipitation

### Is trait variation constrained at higher aridity?

Evidence for increasingly constrained trait variation at higher levels of aridity was mixed, both within and across species. Variance decomposition revealed huge variability in the total amount and dominant scales of within-species trait variation (Figure 4). Variation between plots in a site was almost always the smallest variance component. The relative contribution of within-tree, within-plot and between-site variation differed drastically, however, depending on the trait and species (Figure 4). The only exception was the consistently high amount of within-tree variation in log_10_-transformed HV, which made up >40% of total trait variation in all species. *Acacia acuminata* also tended to have much larger intra-specific trait variation than any of the sampled eucalypts.

**Figure 4:**
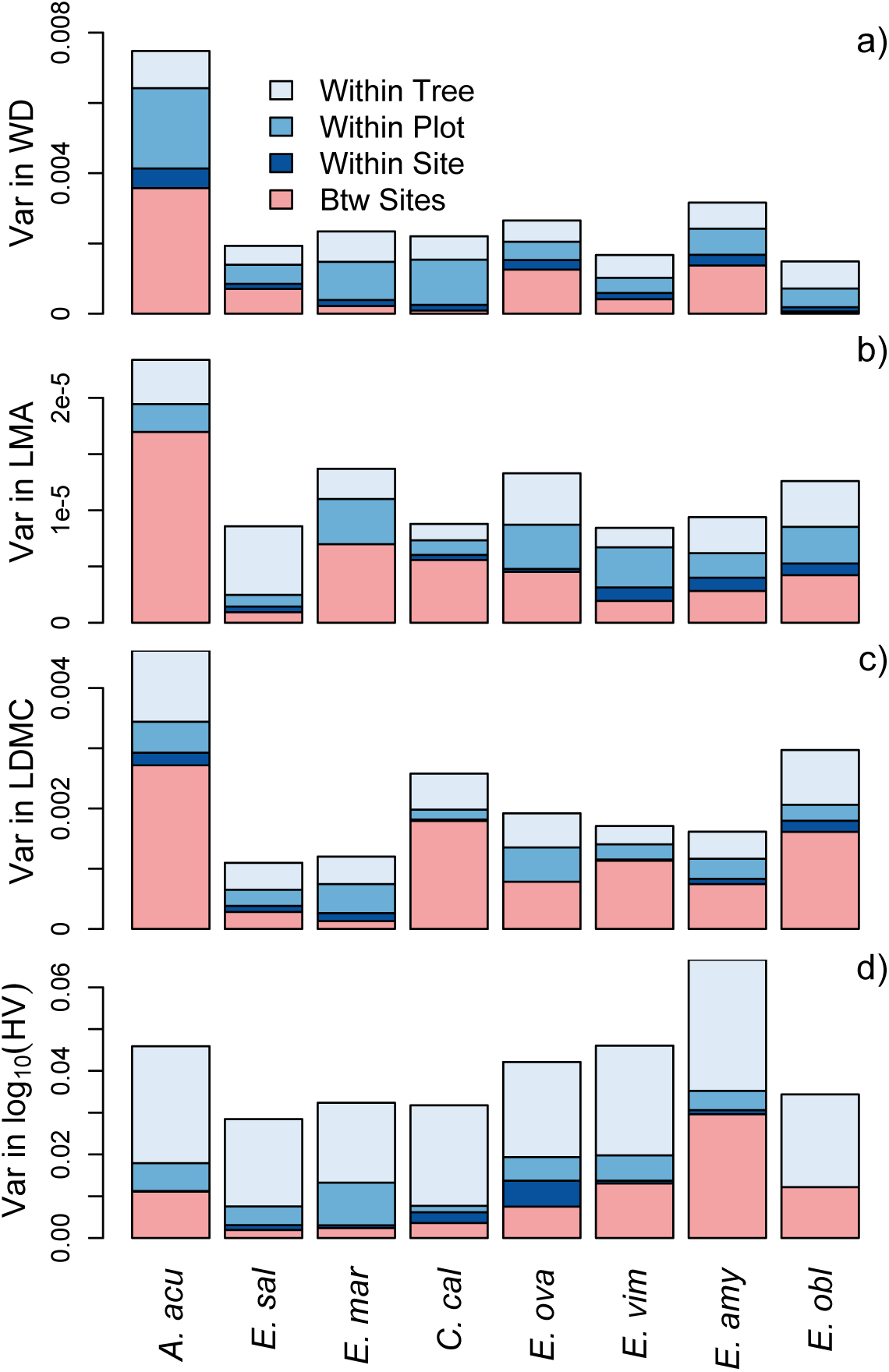
Variance decomposition of four leaf and stem traits measured across the aridity range of eight tree species. The amount and dominant scale of trait variance differs considerably between species for the same trait and between traits. However, variation between plots at a site was almost universally the smallest variance component for all traits and species. Within-tree variation was also always larger for log_10_-transformed A_l_:A_s_ than for all other traits. Species are ordered from driest on the left to wettest on the right. Species abbreviations as in Fig. 1.

Across species, there was limited evidence for decreased intraspecific trait variation in more xeric species. In the seven eucalypts, total within-species trait variation was unrelated to the aridity of species’ driest range edge (the 90^th^ percentile MD of herbarium specimen locations) for WD, LMA, and HV, but was marginally negatively related for LDMC (p=0.092; Figure 5a-d). Most individual variance components were also unrelated to species aridity niche. However, the amount of between-site variation was negatively related to species aridity niche for both LDMC and HV (p<0.01; Figure 5e-h). Results were the same but more statistically significant when using trait coefficients of variation (CV=trait standard deviation divided by trait mean) rather than trait variances (Figure S4).

**Figure 5:**
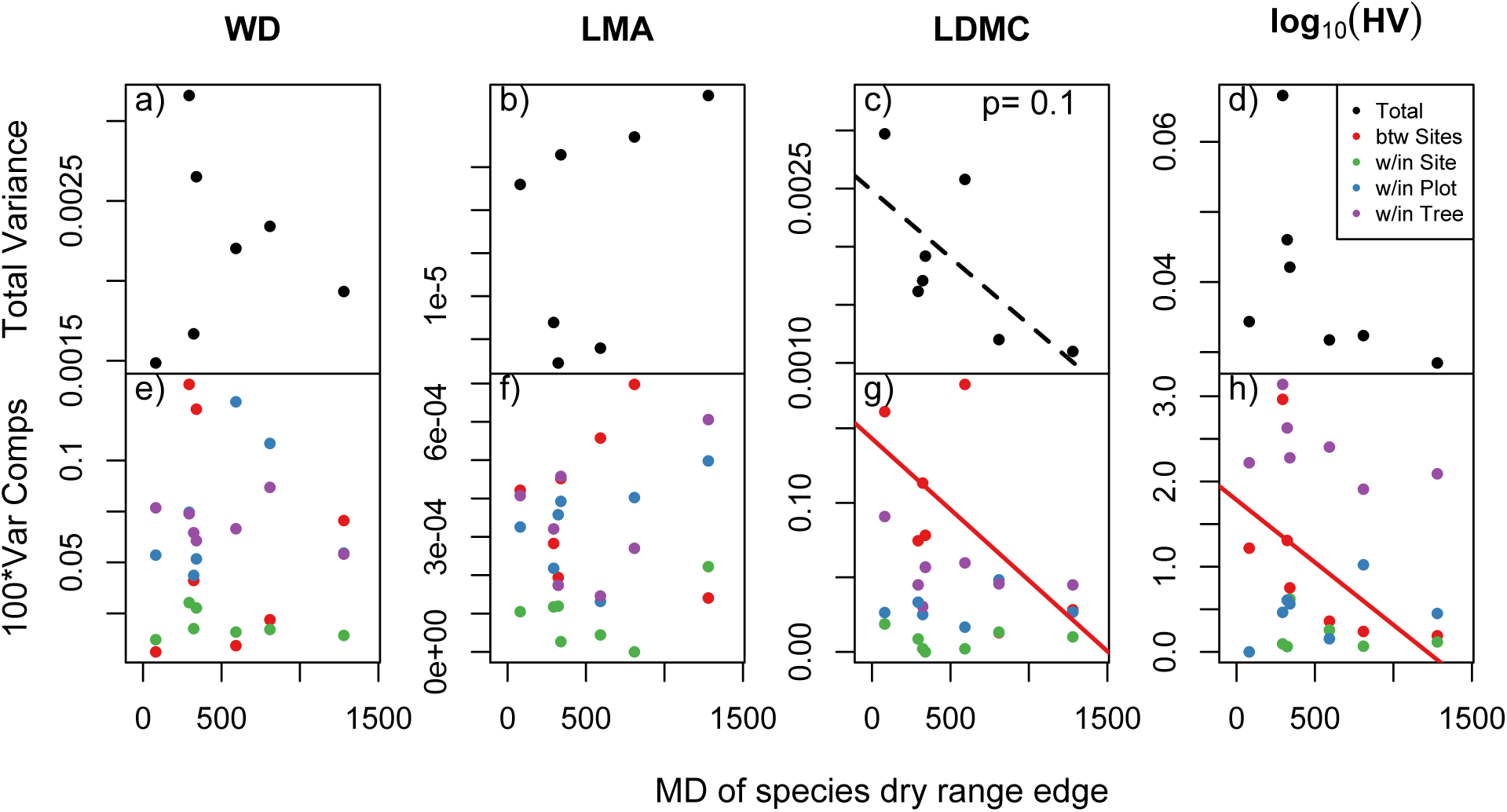
The total amount of within-species trait variation (top row, excluding *A. acuminata*) and potentially climate-related trait variation (variation between sites, red in bottom row), were rarely related to species aridity niche (here shown as the moisture deficit, or mean annual potential evapotranspiration minus mean annual precipitation, of each species driest 90^th^ percentile distribution based on occurrence records in the Atlas of Living Australia). Total trait variation in LDMC decreased marginally significantly in drier species, and climate-related (between site) trait variation in LDMC and A_L_:A_S_ decreased significantly in drier species, consistent with environmental filtering limiting constraining trait variation.

Within-species, variance patterns moving from wet sites to dry sites also showed mixed support for decreasing variance with increasing aridity. A few species did show constrained within-tree and within-plot trait variation at drier sites in a few traits (e.g. LMA in *E. ovata*; Figure 6a), consistent with an increasingly strong environmental filter. Most species for most traits showed no change in trait variance across sites (Figure 6b). LDMC showed the most consistent variance constraint with aridity, with three of eight species showing lower trait variances at drier sites. Even for LDMC, however, almost an equal number of species (two of eight) showed *increasing* trait variances at drier sites. HV showed no aridity-related variance patterns in any species (Figure 6).

**Figure 6:**
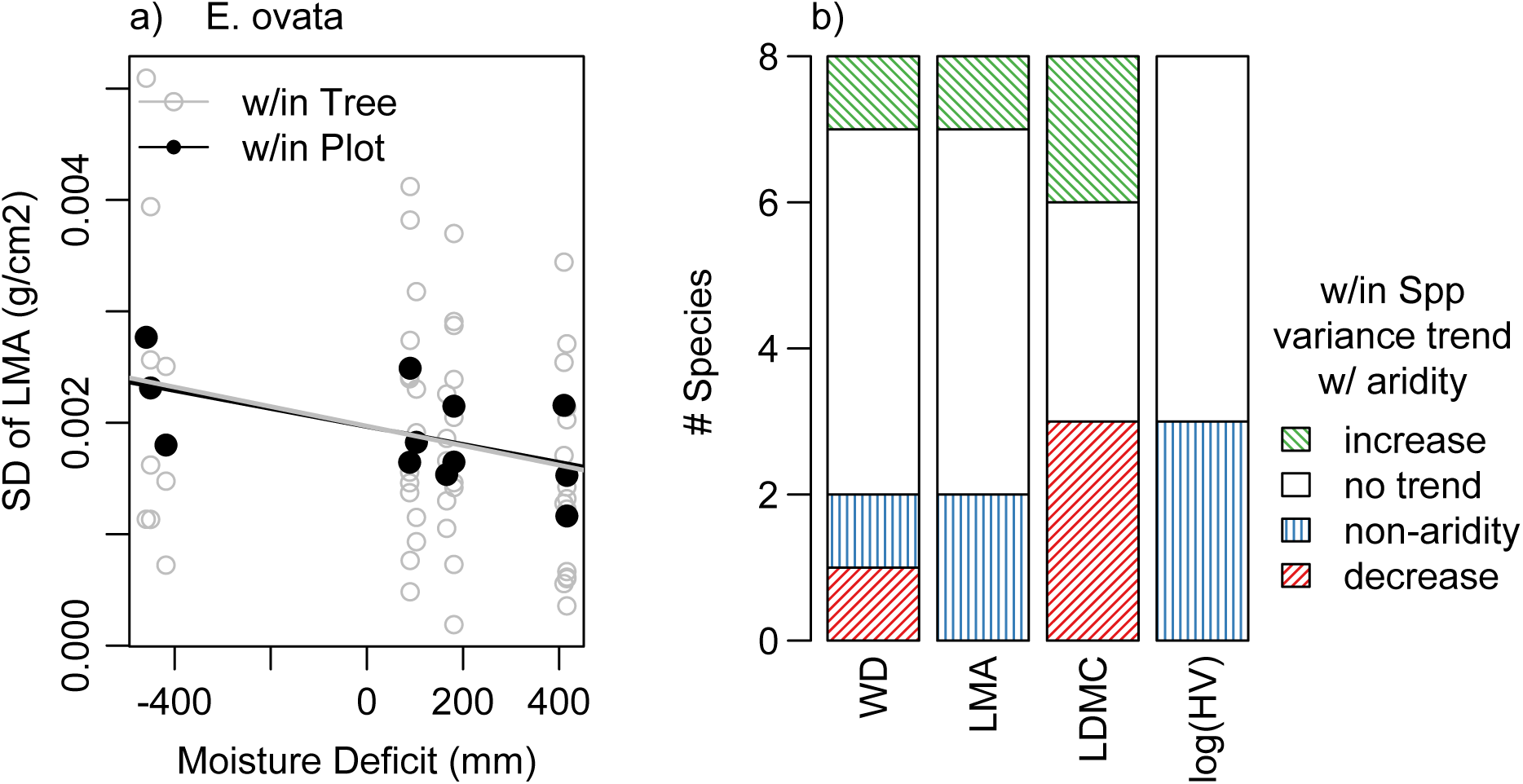
(a) The standard deviation of LMA within tree canopies (grey) and within plots (black) decreases in arid sites (low mean annual precipitation, PPT) in *Eucalyptus ovata*, consistent with increased selection for high LMA values in dry sites. (b) However, overall few species showed evidence of decreasing trait variation (red) at dryer sites, with even LDMC (the trait in which this pattern is most prevalent) showing decreases in only 38% of species and increases in 25% of species. In (b) “non-aridity” signifies species that showed significant site-to-site differences in trait variance that could not be explained by site aridity.

## Discussion

In all, our extensive dataset of 1620 paired trait measurements demonstrated that increasing aridity did indeed result in coordinated trait shifts towards more robust leaves and stems and more hydraulic support per unit leaf area. However, these coordinated trait shifts were only evident across large aridity gradients, implying perhaps indirect mechanistic links between traits and important roles for non-climatic drivers of intraspecific trait variation. Moreover, unlike the consistent patterns in trait means, we did not find decreasing trait variances with decreasing water availability, implying that neither the ‘environmental filter’ nor directional selection seems to consistently limit variation of these particular traits in populations or species living in the driest environments. Below, we discuss these results in greater detail.

### Mean trait shifts

Shifts in leaf, stem, and allocation traits towards more drought resistant values at drier sites were ubiquitous across the sampled species (Figure 1), as might be predicted from correlations between tissue robustness and tissue drought tolerance (e.g. between WD and xylem resistance to embolism; (Chave *et al.* 2009). Indeed, all four traits have previously been reported to show within-species patterns related to water availability either geographically (Schulze *et al.* 2006; Martinez-Vilalta *et al.* 2009; Laforest-Lapointe *et al.* 2014; Niinemets 2014) or experimentally (Poorter *et al.* 2009; Baird *et al.* 2017). However, there are numerous instances where some or all of these traits do not show significant within-species geographic variation related to aridity (Martinez-Vilalta *et al.* 2009; Fajardo & Piper 2010; Richardson *et al.* 2013b; Laforest-Lapointe *et al.* 2014; Vilà-Cabrera *et al.* 2015; Anderegg & HilleRisLambers 2015; Rosas *et al.* 2019). This may be in part due to the nature of these morphological traits themselves. HV is directly relevant to the water balance and hydraulic status of a plant (Whitehead & Jarvis 1981; Trugman *et al.* 2019), but traits like wood density are only partially mechanistically linked to more drought-relevant physiological traits such as xylem vulnerability to embolism (Lens *et al.* 2010). However, complicated and inconsistent trait-environment relationships are often found even for more labor intensive plant hydraulic traits (Rosas *et al.* 2019).

We suggest that the lack of consistency of trait-by-environment relationships in the literature, even for the same species in different studies (e.g. *Pinus sylvestris* in (Martinez-Vilalta *et al.* 2009) vs (Laforest-Lapointe *et al.* 2014)) might be due in part to differences in the extent of the species aridity niche sampled in previous studies, and the presence of confounding environmental gradients in the field. In the literature, it is far more common to find trait-by-aridity relationships in only one of leaf robustness, stem robustness, or leaf vs stem allocation than in all traits, simultaneously. Moreover, studies sometimes uncover apparently contrasting strategies of adjusting robustness versus allocation in different species (Anderegg & HilleRisLambers 2015). This might be due to fundamentally different capacities of various species, genera or clades to adjust tissue characteristics. Even though we found similar patterns in a co-occuring *Acacia*, it is possible that eucalypts are an anomalous taxon. Indeed, previous investigations of Specific Leaf Area (leaf area per unit mass, or the inverse of LMA) found consistent within-species shifts towards more robust leaves at higher aridity across a large number of eucalypt species and multiple, vast aridity gradients (Schulze *et al.* 1998; 2006), which could indicate that eucalypts as a clade are uniquely morphologically flexible. However, other methodological causes of the discrepancies in the literature warrant mentioning.

This study was unique in that it explicitly sampled as much of each focal species’ entire geographic aridity niche as possible, and because the aridity gradients in Australia are largely unconfounded by other stress gradients, notably freezing stress. Given that between-site, or climate-related trait variation is often less than half of total within-species trait variation (Figure 4), sampling as broad of climate space as possible may be critical to ensure that one can detect the climate signal from the considerable noise. Additionally, morphological traits such as LMA are known to vary with multiple environmental signals, including water availability, nutrient availability, and cold stress (Poorter *et al.* 2009). In our study, none of our sites experienced significant cold stress, though soil quality and water availability unsurprisingly co-varied (Table S1, Figure S2). While this means that some fraction of the patterns found here may be due to changes in nutrient rather than water availability (soil quality or depth was the best trait predictor in ∼1/3 of trait-environment relationships), these stresses tend to have similar effects on morphology that may reinforce each other in our study (Poorter *et al.* 2009). However, in temperate study systems, cold stress and water availability tend to have the similar effect of increasing tissue robustness but are often *negatively* correlated on the landscape. We posit that other studies, particularly studies focused on elevational gradients (Fajardo & Piper 2010; Anderegg & HilleRisLambers 2015) and latitudinal gradients (Martinez-Vilalta *et al.* 2009) are likely to see confounding effects of cold stress and drought stress, particularly on leaf traits (González-Zurdo *et al.* 2016; Niinemets 2016). If stem versus leaf allocation (i.e. HV) is less sensitive to cold stress than other morphological traits, this could explain why within-species HV changes across aridity tend to be more ubiquitous than other morphological adjustments (Rosas *et al.* 2019). Thus, the lack of confounding cold stress may explain why we were able to detect aridity-related variation in leaf, stem, and allocation traits in each of seven eucalypts and an *Acacia* (Figure 1, Table S1) with a consistency not previously seen in the literature.

### Trait coordination

We found that coordination across leaf, stem, and allocation traits related to aridity was consistent across species and the dominant mode of trait variation in our study (Figure 2 & 3). One implication of this trait coordination is that the effects of water stress are scaled to species physiology, such that both mesic and xeric species must respond similarly to increasing water stress at their dry range edge regardless of large differences in total water availability. Our seven eucalypt species differed in the wetness of their dry range boundary by over 1000 mm of moisture deficit (Figure 4). Yet all of them showed significant trait-by-aridity relationships and trait-trait coordination across tissues. Even where traits showed weak coordination, namely between branch WD and HV, there was no biogeographic pattern to which species showed strong versus weak trait integration. Both the wettest (*E. obliqua*) and the driest (*E. salmonophloia*) species showed non-significant correlations between WD and HV.

The consistent trait integration across leaf, stem, and allocation traits found here is also reasonably unique in the literature. It is far more common for within-species trait integration to show variable and often unexpected patterns (Richardson *et al.* 2013a; Laforest-Lapointe *et al.* 2014; Anderegg *et al.* 2018; Messier *et al.* 2018; Rosas *et al.* 2019). However, while present in our entire dataset (Figure 3), trait integration only emerged at the largest of spatial and ecological scales (Figure 2g). With the exception of LMA and LDMC, two leaf traits that are highly related both biophysically and physiologically, trait-trait relationships typically only emerged across sites, even though the majority of total trait variation often existed at smaller scales (Figure 4). Indeed, variation between branches in a canopy and between closely located (and likely related) individuals within a plot constituted the majority of trait variation in the majority of traits and species (25 of 32 species’ traits, Figure 4). Despite this, consistent trait correlations, only emerged across site-level trait averages in five of six trait pairs (Figure S2).

This large-scale trait integration suggests that leaf, stem, and allocation traits are only weakly coordinated within species. The relative independence of these traits highlights again that there are many successful ways to be a plant in any given environment. Even when many axes of variation are held constant by looking only within a species, the potential for compensating trait variation (e.g. between roots and leaves) and the important but ultimately weak relationships of many ‘functional traits’ to either physiological rates or demographic outcomes should make weak trait-trait relationships the norm and strong integration the exception in land plants. Moreover, given that functional traits may respond independently to different environmental stresses (Anderegg *et al.* 2018), it should be no surprise that consistent within-species trait integration has been so elusive in the literature. This weak trait integration also highlights that species possess numerous avenues for adapting or acclimating to shifting climate stress in a changing climate.

### Patterns in trait variance

In contrast to the ubiquitous patterns in trait means that we found across all traits and species, we found less evidence for consistent patterns in trait variances with aridity. Looking only across the seven eucalypt species, we found that LDMC and HV tended to be more constrained in xeric than mesic species but the same was not true of LMA and WD. This pattern was not statistically significant for total trait variance, but was significant for between-site variance components for both LDMC and HV, suggesting that the component of trait variation controlled by climate was indeed increasingly constrained at low water availability (Figure 5). However, this pattern only sporadically scaled down to populations within species, with less than half of species showing marked variance patterns across sites for any trait, and LDMC and HV showing similarly weak patterns to LMA and WD (Figure 6).

If these traits are under selection in a warming world (which is likely given the consistent trait-by-aridity relationships documented above), these variance patterns may be good news for the adaptive and/or acclimatory potential of the species in this study. The acclimatory potential for HV may be particularly high, given the consistently high within-tree variation in this trait (Figure 4). Meanwhile, depending on the heritability of WD and LMA, the reliably high within-plot variation (Figure 4) and lack of variance-by-aridity relationships for these traits (Figure 5 & 6) may indicate considerable adaptive potential.

It should be noted, however, that a likely explanation for both the weak trait integration and the mixed variance patterns documented here is that selection is not happening on any of these four traits directly, but rather on other underlying traits that cause some fraction of total trait variation. All four of the studied ‘functional traits’ integrate signals from many different anatomical attributes that have a multitude of influences on actual physiological function (Niinemets 1999; Chave *et al.* 2009; Poorter *et al.* 2009; 2011). Thus, it is common for trait variation in different environments to be driven by disparate anatomical changes that have drastically different physiological consequences but result in identical trait values (e.g. Baird *et al.* 2017).

Within eucalypts, our results might indicate a fundamental constraint on the flexibility of the underlying anatomical properties that drive variation in LDMC and HV, the two traits that did show decreased variance in xeric species (Figure 5). However, a considerable amount of the total variation in both of these traits is non-climatic (Figure 4), making it difficult to detect changes in trait variation at the population level (Figure 6). This highlights the importance of understanding the underlying anatomical drivers of variation in the most common morphological ‘functional traits’ employed by functional ecologists (Niinemets 1999; Onoda *et al.* 2017). Even though we documented consistent within-species trait responses to aridity and trait coordination across tissues, these relationships are unlikely to prove mechanistic in the manner necessary for the parameterization of dynamic ‘trait-based’ vegetation models without gaining a greater understanding of the root causes of this trait variation.

## Conclusion

We found consistent and coordinated trait shifts towards increased tissue robustness and hydraulic supply per unit leaf area across the aridity range of seven related eucalypt tree species and one *Acacia* species. These findings are unique in the literature, in part because we were able to explicitly sample complete aridity gradients that were not confounded by cold stress. However, the compound nature of the gross morphological traits we measured resulted in 1) within-species trait coordination that only emerged across the most climatically disparate individuals in a species and 2) fewer consistent patterns in the size of trait variances with aridity than between trait means and aridity. Our findings imply considerable capacity for these species to adapt and/or acclimate to increasing aridity with future climate change thanks to the substantial within-species variation in multiple traits that is significantly related to climate. Our work highlights outstanding questions about the anatomical mechanisms driving functional trait variation within species, as well as the need to disentangle conflicting effects of different environmental constraints (e.g. temperature versus nutrient versus water) on trait variation to develop a multi-scale understanding of plant functional ecology.

## Acknowledgements

We thank H Wauchope, HR Lai, and J Park for field assistance, T Britton for lab assistance, and G Badgley and A Trugman for comments on the analysis and manuscript. This work was supported by a National Geographic Society Young Explorer Grant (to LDLA). This material is also based upon work supported by the National Science Foundation Graduate Research Fellowship Program under Grant No.s DGE-1256082; DDIG-1500837, an NSF international travel allowance through the Graduate Research Opportunities Worldwide and an NSF Postdoctoral Research Fellowship Grant No. DBI-1711243 and a National Oceanic and Atmospheric Administration Climate and Global Change Fellowship (to LDLA). Any opinions, findings and conclusions or recommendations expressed in this material are those of the author (s) and do not necessarily reflect the views of the National Science Foundation.

