## Supplementary material for "Aridity drives coordinated trait shifts but not decreased trait variance across the geographic range of eight Australian trees": Figure S

[4] Wild Hope Collective

[5] School of Biological Sciences, The University of Queensland, Brisbane, Queensland, Australia

[6] Biological Sciences, School of Natural Sciences, University of Tasmania, Hobart, Tasmania, Australia

[7] Department of Biology, University of Washington, Seattle, WA, USA

[8] School of Biological Sciences, The University of Western Australia, Perth, Western Australia, Australia

**<u>This PDF file contains</u>**

### Table S1: Summary of site characteristics

Site average stand, climate and soil nutrients. DBH: diameter at breast height. Stand BA: stand basal area around focal tree measured using forestry wedge prism (only measured in Tasmania). MD: moisture deficit or PET-PPT. PET: potential evapotranspiration. PPT: annual precipitation. MAT: mean annual temperature. T min: minimum temperature of the coldest month. Climate data are derived from the 1960-1990 climate averages of the Global Aridity and PET Database (Zomer *et al.* 2006) and WorldClim (Hijmans *et al.* 2005). NTO, total soil nitrogen by mass. PTO, total soil phosphorous by mass. Soil data were extracted from the Soil and Landscape Grid of Australia {Grundy:2015dj} and averaged over the top 60cm of the soil column.

| Species | Site Code | Lat (dd) | Lon (dd) | DBH (cm) | Stand BA (ft2 acre-1) | MD (mm) | PET (mm) | PPT (mm) | MAT (°C) | T min (°C) | NTO (%) | PTO (%) | Elev (m) |
|---|---|---|---|---|---|---|---|---|---|---|---|---|---|
| *Acacia acuminata* | BEN | -32.3929 | 118.3881 | NA | NA | 1128 | 1474 | 346 | 16.6 | 4.5 | 0.042 | 0.01 | 356 |
| | GNO | -34.0004 | 118.0661 | NA | NA | 924 | 1287 | 363 | 15.6 | 5.8 | 0.044 | 0.012 | 251 |
| | NAR | -32.982 | 117.08 | NA | NA | 871 | 1363 | 491 | 15.6 | 5 | 0.051 | 0.013 | 356 |
| | PER | -29.4797 | 116.2002 | NA | NA | 1326 | 1654 | 328 | 19.5 | 6.4 | 0.035 | 0.014 | 308 |
| *Corymbia calophylla* | GRIM | -33.6817 | 116.0296 | 40.8 | NA | 422 | 1315 | 893 | 14.7 | 4.6 | 0.099 | 0.014 | 277 |
| | LP | -33.2814 | 116.3329 | 32.9 | NA | 571 | 1360 | 789 | 14.9 | 4.2 | 0.071 | 0.012 | 264 |
| | MIDG | -32.2427 | 116.1469 | 33.6 | NA | 213 | 1331 | 1118 | 16.2 | 5.9 | 0.049 | 0.014 | 334 |
| | STIR | -34.4395 | 118.0727 | 27.1 | NA | 702 | 1203 | 501 | 15.1 | 6.1 | 0.042 | 0.012 | 229 |
| | WAL | -34.9416 | 116.5876 | 37.7 | NA | -69 | 1114 | 1183 | 15.5 | 7.7 | 0.077 | 0.017 | 29 |
| *Eucalyptus marginata* | GRIM | -33.6817 | 116.0296 | 30.6 | NA | 422 | 1315 | 893 | 14.7 | 4.6 | 0.099 | 0.014 | 277 |
| | LP | -33.2814 | 116.3329 | 39.2 | NA | 571 | 1360 | 789 | 14.9 | 4.2 | 0.071 | 0.012 | 264 |
| | MIDG | -32.2427 | 116.1469 | 24.6 | NA | 213 | 1331 | 1118 | 16.2 | 5.9 | 0.049 | 0.014 | 334 |
| | STIR | -34.4395 | 118.0727 | 22.5 | NA | 702 | 1203 | 501 | 15.1 | 6.1 | 0.042 | 0.012 | 229 |
| | WAL | -34.9416 | 116.5876 | 32.4 | NA | -69 | 1114 | 1183 | 15.5 | 7.7 | 0.077 | 0.017 | 29 |
| *E. salmonophloia* | BEN | -32.4056 | 118.3841 | NA | NA | 1128 | 1473 | 345 | 16.7 | 4.5 | 0.042 | 0.01 | 348 |
| | GNO | -33.9632 | 118.0846 | 49.9 | NA | 932 | 1293 | 360 | 15.6 | 5.7 | 0.043 | 0.011 | 260 |
| | NAR | -32.8359 | 117.3283 | 61.3 | NA | 960 | 1380 | 420 | 15.9 | 5 | 0.051 | 0.01 | 357 |
| | PER | -29.4716 | 116.2109 | NA | NA | 1324 | 1654 | 330 | 19.4 | 6.2 | 0.037 | 0.015 | 323 |
| *E. amygdalina* | CMP | -42.9678 | 147.7083 | 42.5 | 57.3 | 167 | 866 | 699 | 12.4 | 4.8 | 0.105 | 0.021 | 34 |
| | EPF | -41.7761 | 147.3177 | 30.5 | 87.3 | 410 | 998 | 588 | 11.4 | 1.8 | 0.086 | 0.016 | 212 |
| | GRA | -42.6287 | 147.4699 | 32.2 | 83.3 | 321 | 934 | 613 | 11.7 | 2.9 | 0.089 | 0.015 | 160 |
| | MER | -41.5616 | 146.2503 | 37.4 | 96.7 | -622 | 852 | 1474 | 9.4 | 1.3 | 0.179 | 0.038 | 471 |
| | TMP | -43.0488 | 147.8975 | 36.2 | 75.3 | -146 | 807 | 954 | 11.9 | 4.8 | 0.127 | 0.027 | 128 |
| *E. obliqua* | BOF | -41.1879 | 148.251 | 60.2 | 120.7 | 61 | 935 | 874 | 12.9 | 4.4 | 0.121 | 0.022 | 56 |
| | GRA | -42.5353 | 147.459 | 44.8 | 101.3 | 235 | 910 | 675 | 10.6 | 1.9 | 0.084 | 0.016 | 320 |
| | MER | -41.5704 | 146.2432 | 45.7 | 98 | -583 | 865 | 1448 | 9.9 | 1.6 | 0.182 | 0.04 | 415 |
| | TMP | -42.9848 | 147.9154 | 46.4 | 162.7 | -164 | 812 | 975 | 11.6 | 4.4 | 0.136 | 0.025 | 184 |
| *E. ovata* | CMP | -42.97 | 147.7045 | 40.3 | 74 | 176 | 867 | 691 | 12.4 | 4.8 | 0.106 | 0.021 | 27 |
| | EPF | -41.7745 | 147.317 | 33.6 | 94.7 | 414 | 999 | 585 | 11.4 | 1.8 | 0.096 | 0.017 | 207 |
| | GRA | -42.4878 | 147.4999 | 32.4 | 71.3 | 94 | 856 | 762 | 9.3 | 0.9 | 0.116 | 0.028 | 539 |
| | MER | -41.5582 | 146.3332 | 29.6 | 76.7 | -442 | 879 | 1321 | 10 | 1.6 | 0.206 | 0.045 | 385 |

| | | | | | | | | | | | | | |
|---|---|---|---|---|---|---|---|---|---|---|---|---|---|
| *E. viminalis* | EPF | -41.776 | 147.3181 | 34 | 98 | 412 | 999 | 587 | 11.4 | 1.8 | 0.096 | 0.017 | 210 |
| | FREY | -41.9583 | 148.137 | 35.8 | 89.3 | 202 | 932 | 730 | 12.1 | 3.5 | 0.135 | 0.019 | 155 |
| | GRA | -42.6287 | 147.4699 | 39.9 | 65.3 | 321 | 934 | 613 | 11.7 | 2.9 | 0.089 | 0.015 | 160 |
| | MER | -41.5782 | 146.2725 | 41.3 | 78.7 | -552 | 867 | 1419 | 9.9 | 1.5 | 0.176 | 0.04 | 423 |
| | TMP | -43.032 | 147.9333 | 17.3 | 153.3 | -168 | 807 | 975 | 11.7 | 4.7 | 0.123 | 0.031 | 143 |

### Table S2: Summaries of Trait-by-environment relationships

Best environmental predictor of within-species variation in mean leaf and stem traits. 'Environ' indicates the climate or soil variable included in the most parsimonious model based on AIC-based model select, and 'p-value' indicates the significance of a Likelihood Ratio Test of the most parsimonious model compared to a null model including no environmental variables a climate variable is list. 'Var pattern' indicates whether there was statistically significant evidence for differing within-population trait variances ('increase' = variance increases with aridity, 'decrease' = variance decreases with aridity, 'non-drought' = significantly different variances per site, but not linked to site aridity, and 'none' = no evidence of differing variances per site). PPT= mean annual precipitation, PET = mean annual potential evapotranspiration, 'MD'= mean annual moisture deficit (PET-PPT).

| | | Wood Density | | | LDMC | | |
|---|---|---|---|---|---|---|---|
| **Species** | **Transect** | **Environ** | **p-value** | **Var pattern** | **Environ** | **p-value** | **Var pattern** |
| *Acacia acuminata* | WA | PET | <0.001 | non-drought | MD | <0.001 | decrease |
| *Eucalyptus salmonophloia* | WA | MD | <0.001 | none | PET | <0.001 | increase |
| *E. marginata* | WA | Depth (PPT) | 0.003 (0.01) | none | PET | 0.02 | increase |
| *Corymbia calophylla* | WA | PET | 0.04 | increase | Fert (none) | 0.008 (n.s.) | none |
| *E. ovata* | TAS | Fert (PPT) | <0.001 (0.003) | none | Fert (PPT) | <0.001 (<0.001) | decrease |
| *E. viminalis* | TAS | Fert (PPT) | <0.001 (0.005) | none | PPT | <0.001 | none |
| *E. amygdalina* | TAS | PPT | <0.001 | none | PPT | <0.001 | none |
| *E. obliqua* | TAS | PET | 0.02 | decrease | MD | <0.001 | decrease |

| | | LMA | | | log(HV) | | |
|---|---|---|---|---|---|---|---|
| **Species** | **Transect** | **Environ** | **p-value** | **Var pattern** | **Environ** | **p-value** | **Var pattern** |
| *Acacia acuminata* | WA | MD | <0.001 | none | MD | 0.031 | non-drought |
| *Eucalyptus salmonophloia* | WA | PET | 0.013 | increase | MD | 0.028 | none |
| *E. marginata* | WA | PPT | 0.008 | non-drought | none | n.s. | non-drought |
| *Corymbia calophylla* | WA | Fert (PPT) | 0.003 (0.048) | none | Depth (PPT) | 0.004 (0.005) | none |
| *E. ovata* | TAS | Fert (PPT) | <0.001 (<0.001) | none | PPT | 0.001 | none |
| *E. viminalis* | TAS | MD | 0.003 | none | Fert (PPT) | <0.001 (0.007) | none |
| *E. amygdalina* | TAS | PPT | <0.001 | non-drought | PPT | <0.001 | non-drought |
| *E. obliqua* | TAS | PET | 0.001 | none | Depth (PPT) | 0.002 (0.042) | none |

### Table S3: Summary of trait-trait correlations at different nested scales

Mean (interquartile range) of Pearson correlations between trait pairs assessed at different scales of organization, summarized across all species.

| trait_pair | Btw site means | Btw plot means in site | Btw tree means in site | Btw branch means in plot |
|---|---|---|---|---|
| **LDMC -vs- LMA** | **0.73 (0.74 - 0.97)** | **0.52 (0.42 - 0.94)** | **0.49 (0.4 - 0.72)** | **0.52 (0.36 - 0.77)** |
| **log.HV -vs- LDMC** | **0.61 (0.51 - 0.84)** | 0.06 (-0.76 - 0.88) | 0.11 (-0.09 - 0.38) | 0.13 (-0.07 - 0.35) |
| **log.HV -vs-LMA** | **0.7 (0.65 - 0.9)** | 0.15 (-0.57 - 0.77) | **0.28 (0.06 - 0.58)** | **0.25 (0.04 - 0.46)** |
| **log.HV -vs-WD** | **0.66 (0.54 - 0.88)** | 0.08 (-0.6 - 0.77) | -0.01 (-0.2 - 0.16) | -0.01 (-0.23 - 0.2) |
| **WD -vs- LDMC** | **0.67 (0.72 - 0.92)** | 0.2 (-0.5 - 0.8) | 0.17 (-0.08 - 0.4) | 0.09 (-0.12 - 0.28) |
| **WD -vs- LMA** | **0.76 (0.69 - 0.87)** | 0.05 (-0.74 - 0.92) | 0.06 (-0.17 - 0.32) | 0.09 (-0.14 - 0.3) |

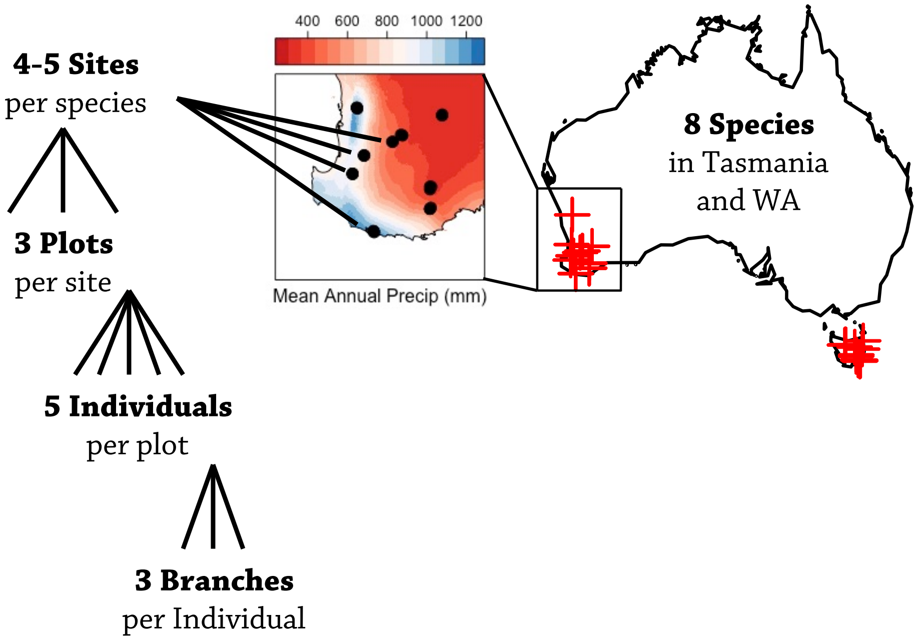

### Figure S1: Diagram of sampling strategy

Red '+' on map of Australia and black points on inset of southern Western Australia indicate sampling sites.

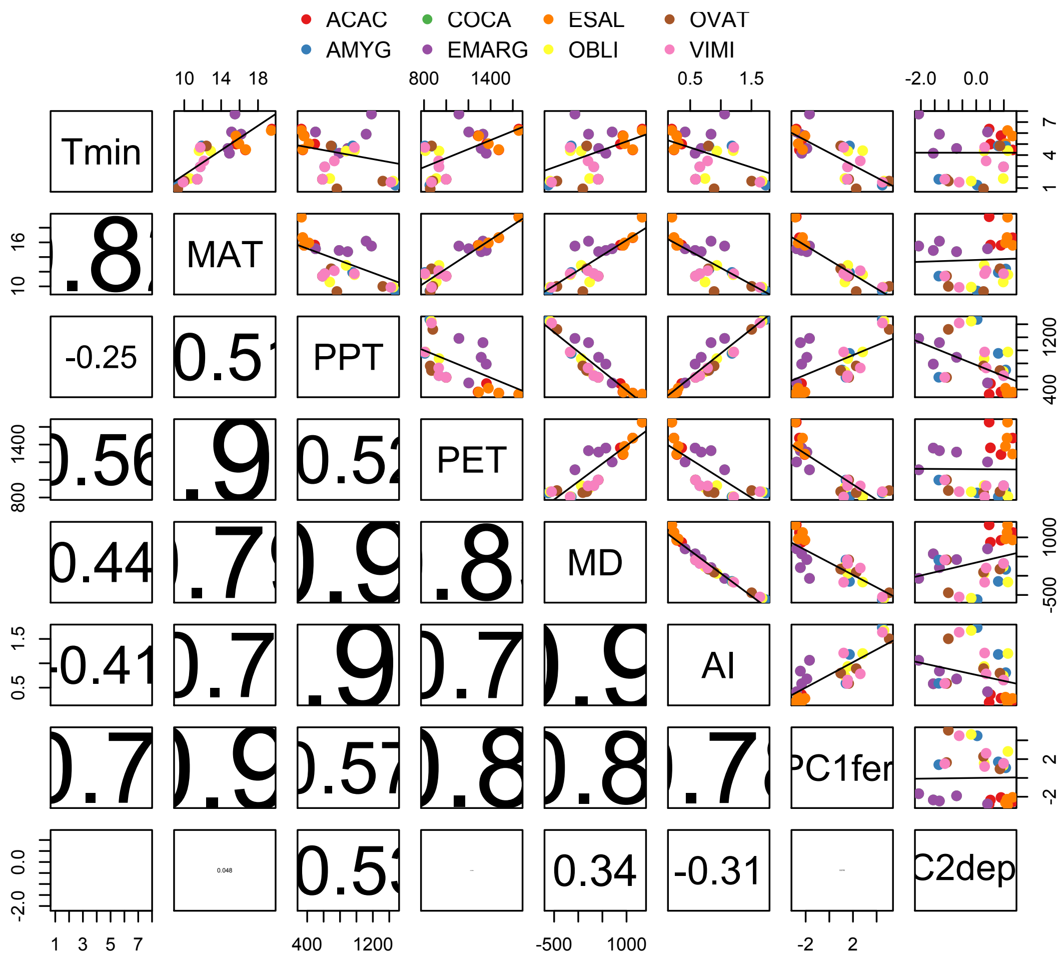

### Figure S2: Correlation between environmental variables across sites

Scatterplot (upper triangular) and correlations coefficient (lower triangular) pairplots between climatic and soil variables. Correlation coefficients are scaled in size by their absolute magnitutde. Tmin = minimum temperature of coldest month, MAT = mean annual temperature, PPT = annual precipitation, PET = potential evapotranspiration, MD = moisture deficit (PET – PPT), AI = aridity index (PPT/PET), PC1fert = soil PC1 representing soil fertility, PC2depth = soil PC2 representing soil depth. Species abreviations: ACAC – *Acacia acuminata,* AMYG – *E. amygdalina,* COCA – *Corymbia calophylla,* EMARG – *E. marginata,* ESAL – *Eucalyptus salmonophloia,* OBLI – *E. obliqua,* OVAT – *E. ovata,* VIMI – *E. viminalis*

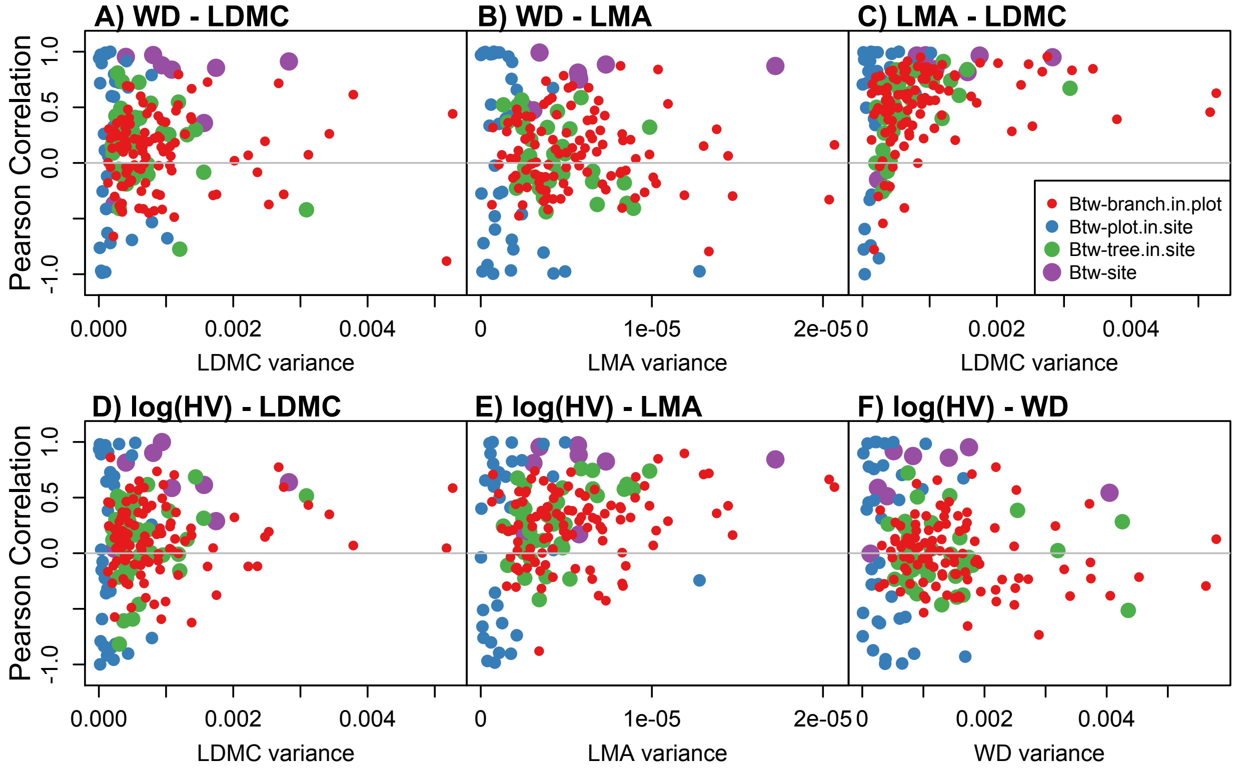

### Figure S3: Funnel Plots of trait covariation

Correlations between site mean trait values were reasonably strong for all eight focal species for all trait pairs, however little trait coordination was observed at lower levels of organization (within plots, within sites) for all trait pairs except for LMA-LDMC. With the exception of the relationship between log(Al:As) and LMA, the lack of trait correlations at lower levels of organization did not appear to be driven by limited trait variances at lower levels (i.e. correlations do not converge towards the purple between-site correlations at higher x-values).

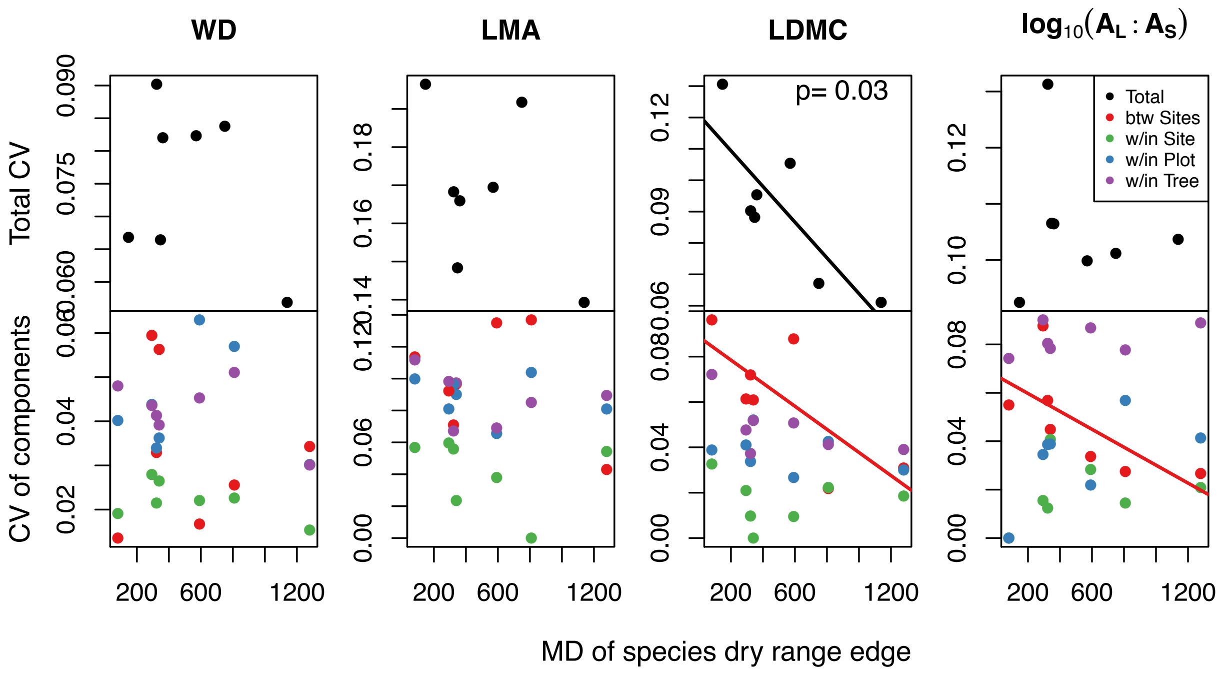

### Figure S4: Trait CVs versus climate niche

The total within-species trait coefficient of variation (CV= trait standard deviation/trait mean, top row) and CV of potentially climate-related trait variation (variation between sites, red in bottom row), were rarely related to species aridity niche (here shown as the moisture deficit, or mean annual potential evapotranspiration minus mean annual precipitation, of each species driest 90$^{th}$ percentile distribution based on occurrence records in the Atlas of Living Australia). Total trait CV of LDMC decreased significantly in drier species, and climate-related trait CV in LDMC and Al:As decreased significantly in drier species, consistent with environmental filtering limiting constraining trait variation.
